# Structural variability of apolipoprotein A-I amyloid fibrils across organs, mutations, and clinical presentations, revealed by cryo-EM

**DOI:** 10.1101/2025.07.08.663734

**Authors:** Binh An Nguyen, Maria del Carmen Fernandez-Ramirez, Parker Bassett, Virender Singh, Preeti Singh, Maja Pękała, Layla Villalon, Yasmin Ahmed, Andrew Lemoff, Bret Evers, Christian Lopez, Barbara Kluve-Beckerman, Lorena Saelices

## Abstract

Hereditary apolipoprotein A-I (AapoA-I) amyloidosis is a rare systemic disease caused by the deposition of amyloid fibrils formed by apolipoprotein A-I in multiple organs, leading to severe clinical outcomes. With no available therapies or diagnostic tools, defining the structure of AApoA-I fibrils is crucial to understanding disease mechanisms and guiding intervention. Using cryo-electron microscopy, we analyzed AApoA-I fibrils from the heart, kidney, liver, and spleen of patients carrying G26R, L90P, and R173P mutations. G26R fibrils, regardless of organ, exhibited untwisted morphologies and could not be resolved structurally. Conversely, L90P and R173P fibrils displayed a compact diabolo-shaped conformation in all organs analyzed. Their high-resolution maps enabled visualization of *cis*-Proline 66, which may represent a potential conformational switch during fibril formation. Our findings suggest that mutation-driven polymorphism may influence organ tropism and clinical presentation. This work advances our understanding of AapoA-I fibril assembly and provides insights toward developing targeted clinical tools.

## Introduction

Hereditary apolipoprotein A-I amyloidosis (AApoA-I amyloidosis) is a rare systemic disease caused by the deposition of amyloid fibrils composed of apolipoprotein A-I (ApoA-I). These fibrils accumulate in multiple organs, including the liver, kidneys, heart, peripheral nerves, gastro-intestinal tract, skin, eyes and testes, leading to tissue damage, organ failure, and eventually death^1^. Disease onset varies widely, with symptoms often arising between the ages of 20 and 70^2^. Standard amyloid testing, such as Congo Red or thioflavin S staining, do not differentiate AApoA-I amyloidosis from other amyloid types without genetic or proteomic analysis, often resulting in misdiagnosis and suboptimal treatment^3^. Current treatment is primarily supportive, with liver transplantation as the main option to replace mutated ApoA-I with a wild-type ApoA-I, or organ transplantation to address failing or damaged organs^4^. Similar to other rare amyloidosis diseases, a major challenge in the field of AApoA-I amyloidosis is the limited understanding of disease mechanisms leading to variability of clinical presentations.

ApoA-I is a multifunctional protein with a well-defined structural organization that plays a central role in lipid metabolism and cardiovascular protection. The matured ApoA-I protein consists of 243 residues primarily synthesized in the liver and the intestines^5^. Its native structure is predominantly amphipathic α-helices making up of ten tandem repeats that allow it to interact with phospholipids effectively^6-8^. ApoA-I has three functional domains that contribute to its functions in lipid metabolism. An N-terminal domain (residues 1-184) stabilizes the lipid-free ApoA-I, facilitates binding to lipid due to its amphipathic α-helix structure, and enables interactions with lipids to form of high-density lipoproteins (HDLs)^9^. Overlapping with this domain, a central region (residues 144-164) is crucial for activating lecithin-cholesterol acyltransferase, an enzyme essential for cholesterol esterification within HDL particles, and for cellular cholesterol efflux^10,11,12^. Finally, a C-terminal domain (residues 185-243) allows ApoA-I to anchor to lipid surfaces, increasing its binding capacity^13,14^. Functionally, wild-type ApoA-I is a crucial component of HDLs, which are responsible for removing excess cholesterol from peripheral tissues and transporting it to the liver for excretion. This cardiovascular protective mechanism, known as reverse cholesterol transport, helps mitigate atherosclerosis by reducing lipid buildup in blood vessel walls and maintaining cholesterol balance in the body^10,13^.

Despite its protective role, both wild-type and variant ApoA-I can misfold and aggregate under certain conditions. To date, over 20 pathogenic mutations in ApoA-I have been identified, with many clustering in two residue stretches: residues 26-75 and 154-178^6,15^. Proteomic analysis of *ex-vivo* AApoA-I amyloid fibrils have consistently shown that the fibril core is primarily composed of N-terminal fragments, typically covering the first 83 to 98 residues, regardless of mutation sites^16-18^. The current evidence suggests that mutations destabilize the native conformation of ApoA-I, exposing the N-terminal aggregation-prone regions (APRs) and initiating amyloid formation^19-21^. Yet, the precise molecular mechanism linking mutations to fibril formation, and how these changes translate to clinical outcomes remains poorly understood.

Previous studies have reported that the position of the ApoA-I mutation correlates with distinct patterns of tissue deposition and subsequent clinical manifestations^22,23^. Specifically, mutations between residues 26 and 75 are associated with deposition in the kidney and liver, while those between residues 90 and 178 tend to localize in the heart, larynx, and skin (Table S1)^22-24^. Additionally, amyloid deposition of wild-type ApoA-I has been observed in the arterial walls of individuals with atherosclerosis^25^. The mechanism by which ApoA-I variant position drives organ tropism remains poorly understood. In neurodegenerative amyloid diseases, such as tauopathies or synucleinopathies, each clinical syndrome is associated with a defined structure (or structures) of the amyloid fibrils^26-28^. Inspired by these studies, we hypothesize that the clinical presentations in AApoA-I amyloidosis may be linked to the structural conformation of the amyloid fibrils, which could, in turn, be determined by the site of the mutation.

In this study, we determined the structures of *ex-vivo* AApoA-I amyloid fibrils extracted from multiple organs and patients. Samples included fibrils extracted from heart, liver, kidney, and/or spleen from three patients: one patient with renal and hepatic involvement carrying the G26R mutation, and two with cardiac and cutaneous manifestations carrying the L90P and R173P mutations^29-31^. Mass spectrometry and cryogenic electron microscopy (cryo-EM) revealed distinct fibril conformations associated with different clinical phenotypes, while structural consistency was observed across tissues within the same patient. Our preliminary data suggests that the mutation site may influence fibril architecture, potentially contributing to phenotypic heterogeneity.

## Results

### Histological study of amyloid deposits

We obtained freshly frozen samples from three patients with hereditary AApoA-I amyloidosis and conducted histological analysis using Congo red, Thioflavin S, and hematoxylin and eosin stains (Fig. 1 and Fig. S1a,b). Due to sample availability, we histologically analyzed only select organs for each patient. However, we identified amyloid deposits in all tissues available for analysis, though the amount of fibril deposition varied. Patient 1 (G26R), a 65-year-old female who died of renal failure, carried the G26R mutation^29^. We observed amyloid deposits in the liver (among hepatocytes, Fig. 1a-1), spleen (in red pulp regions, Fig. 1a-2) and kidney (in the medulla, Fig. 1a-3). Patient 2 (L90P), a 60-year-old female who died of heart failure and presented cutaneous lesions, carried the L90P mutation^31^. We observed amyloid deposits around the epicardial coronary artery of the heart (Fig. 1b). Patient 3 (R173P), a 63-year-old male who died of heart, renal, and liver failure, carried the R173P mutation^30^. We observed amyloid deposits in the liver (among hepatocytes, Fig. 1c-1), spleen (surrounding the central artery, Fig. 1c-2), kidney (in the hilar fascia, Fig. 1c-3), and heart (in the extracellular space of myocardial cells, Fig. 1c-4).

**Figure 1.**
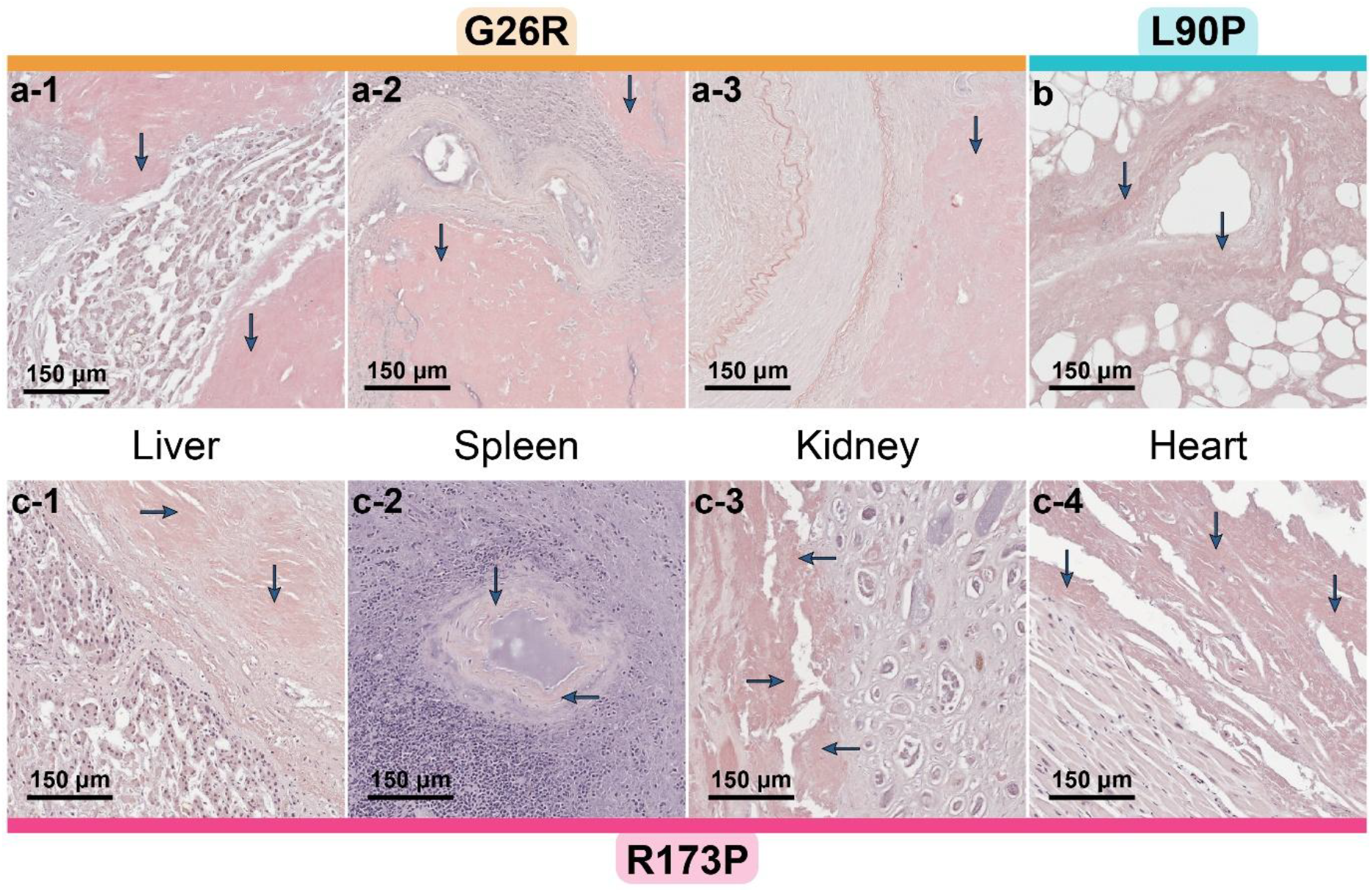
**Histological analysis of AApoA-I amyloid deposits** of various tissue samples from three patients, G26R, L90P and R173P, using Congo red staining. Arrows indicate amyloid deposits. Cross sections show amyloid in the liver, spleen and kidney of G26R (a1-3), the heart of L90P (b), and the liver, spleen, kidney and heart of R173P (c1-4). Scale bar: 150 µm.

### Fibril extraction and protein identification by mass spectrometry

We purified AApoA-I fibrils from all samples using an ice-cold water extraction protocol, as previously described^32^. Transmission electron microscopy (TEM) of the resulting eluates confirmed the presence of amyloid fibrils in all tissue samples (Fig. S2), consistent with histological staining. We confirmed the ApoA1 identity of these isolated *ex vivo* fibrils using mass spectrometry. Tryptic analysis revealed ApoA-I sequence coverage ranging from 71 to 89% across all fibril extracts (Fig. S3a). Using intact mass spectrometry, we identified a fragment spanning residues 1 to 81 as the predominant species across all tissues and patients (Fig. S3b). Additional patient-specific fragments were also identified: fragments spanning residues 1-67 and 1-71 in all G26R samples, 1-85 and 1-93 in all L90P samples, and 1-85 in all R173P samples (Fig. S3b). We confirmed heterozygosity in all patients, consistent with previously reported, as tryptic and GluC digestions generated fragments containing both the wild-type and variant residue at the mutation site (Fig. S3a).

### Cryo-EM reveals structural heterogeneity of G26R fibrils vs both L90P and R173P fibrils

We prepared cryo-EM grids using elution fractions that contained high-quality fibrils, based on fibril concentration and morphology assessed by TEM (Fig. S2). We excluded three extracts— the G26R liver sample and the R173P liver and spleen samples—because of excessive fibril bundling or low yield. The remaining fibril extracts showed well-dispersed, abundant fibrils, which enabled cryo-EM structure determination. These included fibrils from the kidney and spleen of the G26R patient, the heart of the L90P patient, and the heart and kidney of the R173P patient.

Two-dimensional (2D) class averages provided initial insight into the structural characteristics of fibrils across patient samples. (Fig. 2a). Renal and splenic fibrils from the G26R patient displayed diameters of approximately 7.5 to 19 nm and lacked apparent twist, rendering them unsuitable for helical reconstruction (Fig. 2b). In contrast, cardiac fibrils from the L90P and R173P patients, as well as renal fibrils from the R173P patient, exhibited twisted morphology with diameters of approximately 9.5 nm (Fig. 2b and Fig. S4a-b). We used the resulting 2D classes from twisted fibrils to generate initial models for further structural processing.

**Figure 2.**
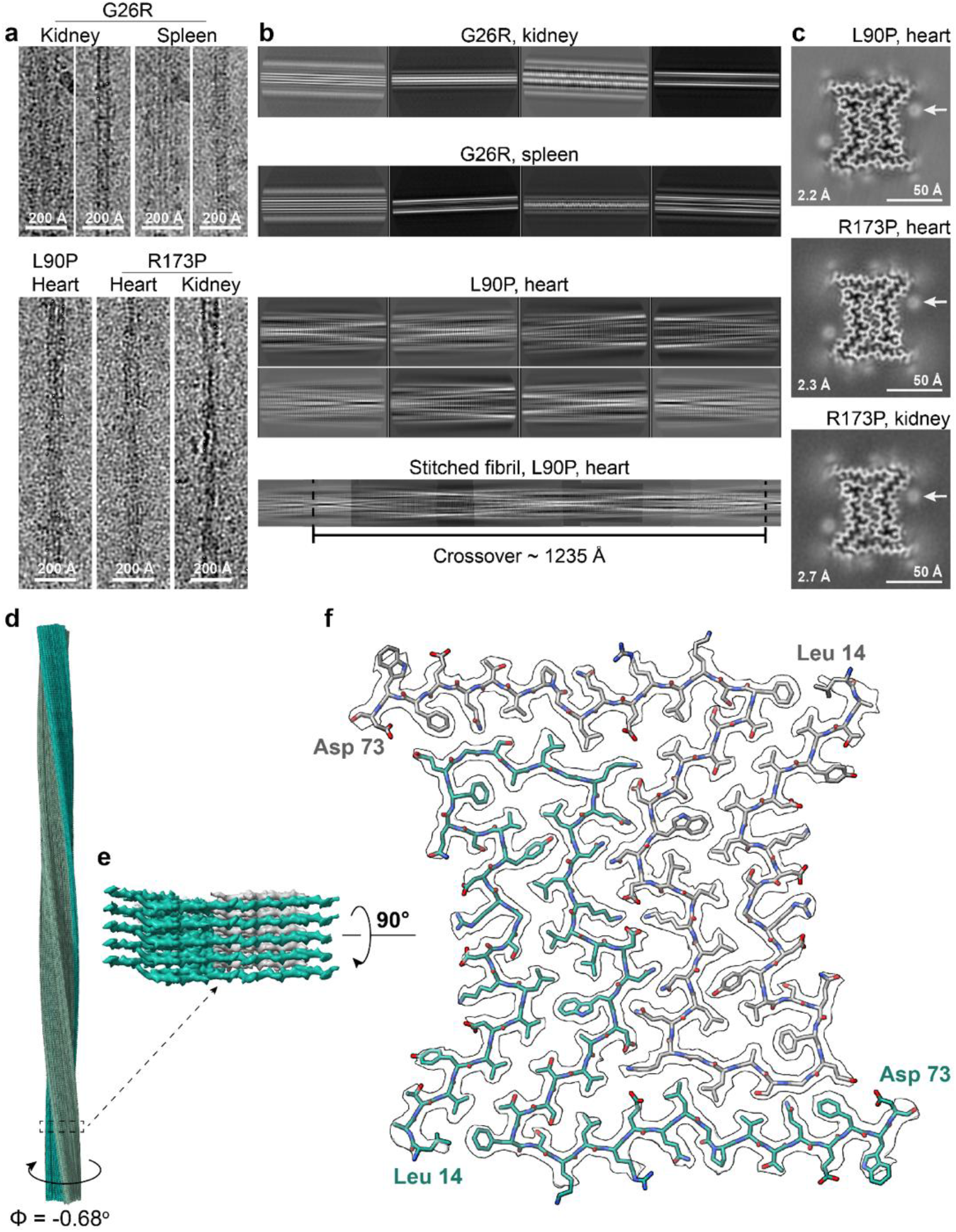
Cryo-EM images of AApoA-I fibrils, their 2D and 3D classifications, and the reconstructed fibril from L90P variant. **(a)** Cryo-EM images of fibrils from G26R kidney and spleen, L90P heart and R173P heart and kidney. Scale bar: 200 Å. **(b)** 2D class averages (particle box size of 300 pixels) of G26R fibrils from the kidney and spleen, and L90P heart. Stitched 2D class averages of fibrils showing the crossover from L90P variant. **(c)** Cross sections through the reconstructed L90P, R173P heart and kidney fibrils, perpendicular to the helical axis. Resolutions in Å are indicated on the bottom left. White arrows point to the extra densities. Scale bar: 50 Å. **(d)** The reconstructed L90P fibril containing 360 layers. **(e)** Magnified view of (d) showing a subset of 5 fibril layers. **(f)** Cryo-EM map and atomic model of AApoA-I fibrils shown for a single layer perpendicular to fibril axis.

We determined the structures of the ordered cores of AApoA-I fibrils from the L90P heart, the R173P heart, and the R173P kidney using helical reconstruction of cryo-EM images, achieving resolutions of 2.2 Å, 2.3 Å, and 2.7 Å, respectively (Fig. 2c and Fig. S5). The fibril folds were identical across all samples, regardless of tissue origin or mutation, indicating structural conservation. Each fibril was composed of two intertwined protofilaments forming a compact, diabolo-shaped core (Fig. 2c). The fibril layers were separated by a rise of 4.75 Å and a twist angle of −0.67° (±0.01) (Fig. 2d,e and Fig. S4c,d). We also observed two unidentified round densities (∼8 Å in diameter) positioned outside the fibril core, along with weaker densities at the fibril ends, suggesting the presence of putative binding molecules and/or disordered regions at both the N- and C-termini of ApoA-I (white arrows in Fig. 2c).

The high resolution of the AApoA-I fibril density maps enabled us to determine the handedness of the helical twist and to build accurate atomic models (Fig. S6a). The ordered core of each protofilament spanned residues Leu 14 or Ala 15 to Asp 73 or Trp 72, placing the mutations outside the structured region (Fig. 2f and. Fig. S4e,f). The alignment of the Cα backbones from the three fibril structures—L90P heart, R173P heart, and R173P kidney— revealed high structural homogeneity, with an average root mean square deviation (r.m.s.d.) of 0.15 Å, as calculated by GESAMT (Fig. S6b-c). The mass spectrometry analysis of fibril extracts supports this model, identifying two major cleavage-prone regions flanking the AApoA-I fibril core in all variants (Fig. S7).

### Analysis of the L90P and R173P fibril structures

AApoA-I fibrils rank among the highly stable amyloid assemblies characterized to date^33^. *In silico* structural analysis estimates their solvation energies at ∼66.6 kcal/mol per layer, −33.3 kcal/mol per molecule, and −0.56 kcal/mol per residue (Fig. S8a). This exceptional stability arises from a dense interaction network that reinforces the protofilament core and supports interprotofilament and interlayer contacts.

Hydrophobic interactions stabilize the protofilament core, while polar interactions maintain the interface between protofilaments. Each AApoA-I molecule contributes six β-strands, plus a loop between Leu22 and Phe33 (Fig. 3a). The first four β-strands form a long hydrophobic hairpin, and β6 interacts with the β2′ strand from the neighboring protofilament (Fig. 3b; *prime indicates the adjacent protofilament*). The two protofilaments also interact through salt bridges and hydrogen bonds, as shown in Fig. S8b and Fig. S9a-b. Additionally, we observe salt bridges on the solvent-exposed surface (Fig. S9c), typical π–π interactions of aromatic residues, hydrogen bonding from stacked aspartate and glutamate side chains and an interlayer hydrogen bonding network formed by the peptide backbone—all characteristic features of the amyloid architecture (Fig. S9d-f). Notably, Proline 66 (n) adopts a cis-isomer conformation in both protofilaments, enabling the formation of a hydrogen bond with Gln 63 from the layer below (n-1) (Fig. S9g). Finally, we observe small densities embedded at the protofilament interface within the fibril, likely water molecules that form hydrogen bonding along the fibril axis (Fig. S9h).

**Figure 3.**
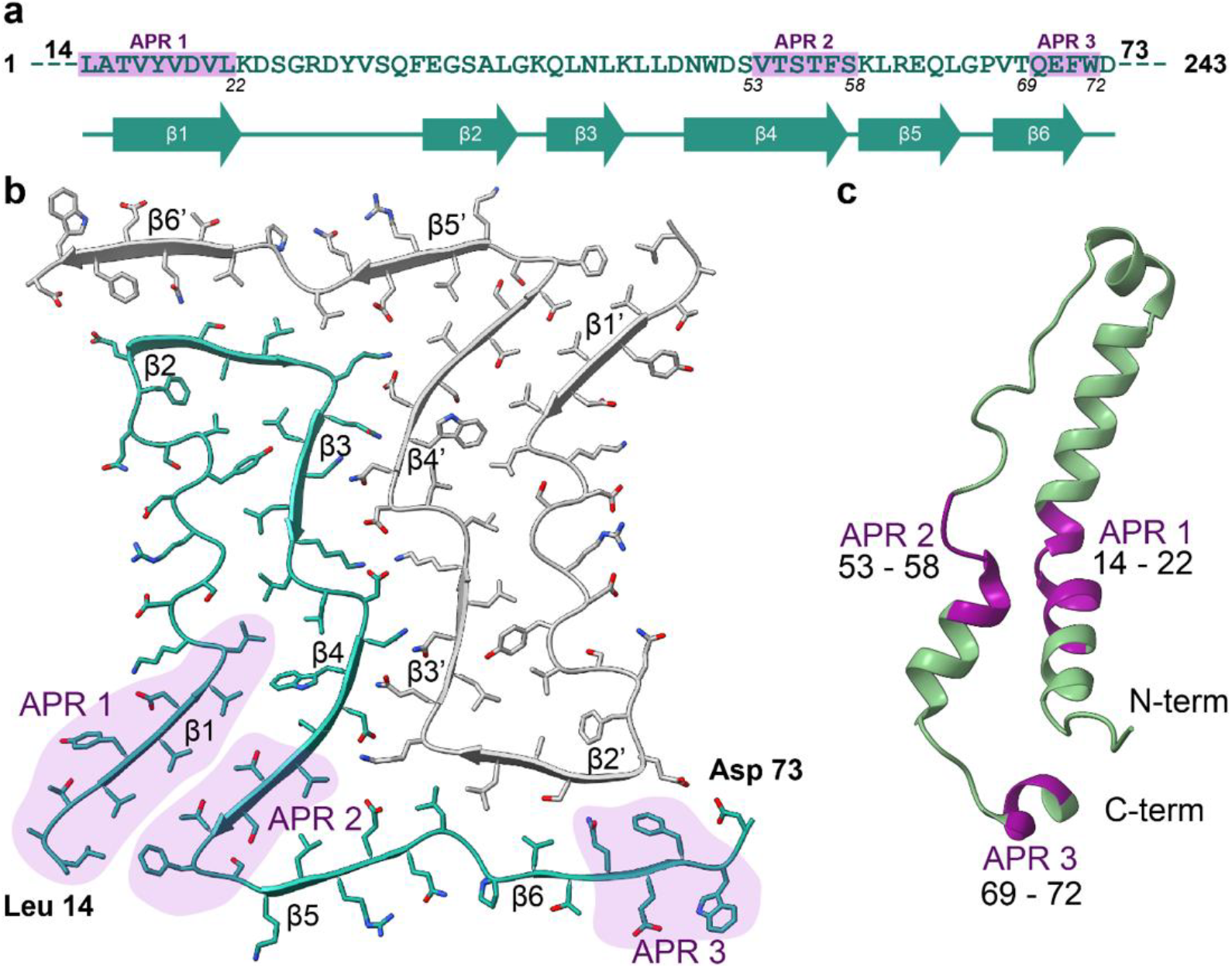
Structural information of AApoA-I fibrils and the location of aggregation-prone regions. **(a)** Amino acid sequence (green) of the AApoA-I fibril core; the regions highlighted in purple are the APRs; green arrows indicate β-strands. **(b)** Top view of a single fibril layer from L90P variant showing the arrangement of β-strands and location of APR1 to APR3 (in purple area). **(c)** Native crystal structure of ApoA-I spanning residues 2-75 (PDB: 3R2P) and the locations of APRs are highlighted in purple.

### Analysis of aggregation-prone regions of AApoA-I

*In silico* analysis of the AApoA-I fibril structure revealed key APRs that potentially drive and stabilize fibril assembly through steric zippers and aromatic interactions. Four key APRs were identified; three located within the AApoA-I fibril, corresponding to β1 (APR1), β4 (APR2) and β6 (APR3) strands (Fig. 3). The β1 and β4 strands form an extended N-to-C hetero-steric zipper that brings APR1 and APR2 together, maintaining a similar proximity as seen in the ApoA-I crystal structure (Fig. 3b,c)^34^. Additionally, the β4 strand establishes a shorter N-to-C zipper on the opposite side, involving interactions between Asn 49 and Asp 51 with Asn 43’ and Gln 41’ on the β3’ strand. The β6 strand formed a N-to-C zipper with the β2’ strand from the adjacent protofilament. Interestingly, all identified APRs within the fibril core include at least one aromatic residue: Phe 33 in APR1, Tyr 50 in APR2, and Phe 71 and Trp 72 in APR3. A fourth APR, spanning residues Val 221–Leu 233 at the C-terminus, was predicted to lie outside the fibril core.

## Discussion

The clinical presentation of AApoA-I amyloidosis varies significantly, including affected organs, symptom severity, and age of onset. Symptoms often overlap with other diseases complicating timely diagnosis and effective clinical management^4,35^. Previous studies have established associations between specific mutation sites and organ involvement, though a mechanism behind this phenomenon has yet to be deduced^24,36^. In this study, we present cryo-EM structures of AApoA-I fibrils isolated from multiple organs of three patients with distinct genotypes and clinical presentations. Our preliminary results suggest that mutations in the *ApoA-I* gene affect the structure of AApoA-I fibrils, which potentially contribute to the observed heterogeneity in disease manifestation.

The location of single-point mutations in ApoA-I has been associated with distinct clinical phenotypes. Studies have shown that mutations within the N-terminal region (residues 1–75) are predominantly linked to renal and hepatic pathologies, while mutations beyond residue 90 are more commonly associated with cardiac and laryngeal involvement, albeit not exclusively^15,36-38^ (Table S1). Our structural analyses demonstrate that fibrils formed by the G26R variant adopt a distinct, untwisted morphology, whereas fibrils carrying L90P and R173P mutations exhibit a consistently twisted morphology (Fig. 2a and b). This morphological divergence suggests that mutation-specific alterations in fibril architecture may play a critical role in modulating tissue tropism and organ-specific amyloid deposition^22^. Comparable correlations between fibril structure and disease phenotype have been observed in tauopathies and synucleinopathies, neurodegenerative diseases characterized by tau and a-synuclein protein aggregation, respectively^26-28^. These diseases include Alzheimer’s, Parkinson’s, chronic traumatic encephalopathy, and Lewy Body dementia, amongst others. Structural studies in these diseases have shown that each clinical syndrome is associated with a unique filament fold, reinforcing the importance of fibril conformation in disease expression. Our analysis of AApoA-I fibrils supports a potential connection between untwisted fibril morphologies and renal/hepatic phenotypes, and twisted fibril structures and cardiac involvement. Further investigation of additional AApoA-I variants across diverse tissues will be critical to elucidate how mutation location influences fibril polymorphism, deposition site, and disease presentation.

To explore the structural basis underlying these phenotypic associations, we examined how mutation location may impact the integrity of the fibril core and, consequently, fibril morphology. Cryo-EM helical reconstruction of L90P and R173P fibrils revealed a conserved diabolo-shaped core composed of two intertwined protofilaments, consistent across multiple organs in these patients (Figs. 2f and S4e,f). This fibril core, spanning residues 14 to 73, appears unaffected by mutations outside this region. In contrast, the G26R substitution introduces a bulky, ionic residue directly into a hydrophobic pocket, potentially disrupting core formation (Fig. S10). Similar structural disruptions may occur with other mutations located within the fibril core, such as W50R, L60R, L64P, and F71Y, due to steric clashes or interference with key hydrophobic interactions (Fig. S10). These destabilizing effects could favor the emergence of alternative fibril morphologies such as the untwisted species observed in the G26R patient (Figure 2a, b).

Mutations may also influence the proteolytic profile of ApoA-I and may be shaped by, or contribute to, the distinct fibril morphologies observed in our samples. *In vitro* studies have shown that mutations such as G26R and L178H increase susceptibility to N-terminal proteolysis compared to wild-type protein, likely due to structural alterations in the lipid-binding C-terminal domain^18,39^. These changes diminish lipid-binding ability, rendering the protein more prone to proteolytic activities^40^. Notably, different mutations can affect lipid binding ability of ApoA-I to varying degrees, leading to divergent proteolytic outcomes^41^. This variability may explain our intact mass spectrometry results, which revealed unique N-terminal fragments of different lengths among fibrils from distinct variants (Fig. S7).

Proteolysis may play a dual role in ApoA-I fibril formation. Some studies propose that proteolysis precedes and facilitates fibril formation by exposing aggregation-prone N-terminal fragments^42,43^. *In vitro* studies demonstrate that full-length ApoA-I, both wild-type and G26R mutant, does not form fibrils under various conditions, whereas the N-terminal fragment (residues 1–83) of both constructs readily assembles into fibrils^43,44^. Alternatively, others studies propose that APRs of ApoA-I become exposed due to mutation-induced conformational changes, with fibril formation preceding proteolysis^19,45^. Another study proposes that full-length ApoA-I may form fibrils upon methionine oxidation *in vitro*^46^. Together, these findings raise the possibility that the unique truncated fragments observed in our study may arise through two scenarios: in one, specific proteolytic fragments seed distinct fibril morphologies; in the other, structural differences in fibrils expose additional protease cleavage sites. Our current data do not allow us to determine the sequence or causality of these events, leaving open questions about the complex relationship between mutation, proteolysis, and fibril formation in AApoA-I amyloidosis.

Our fibril structures offer insights into a potential mechanism of ApoA-I aggregation. While mutation location may dictate fibril structure and proteolysis may be a prerequisite for fibril formation, the exposure of APRs likely drives the process. The pronounced structural difference between the native and amyloid forms of ApoA-I suggests a conformational conversion involving destabilization of the native protein and reassembly into a β-sheet–rich architecture. *In vitro* studies reveal that mutations such as G26R and L178H destabilize the helical structure of full-length ApoA-I, potentially exposing APRs^18,39^. Notably, APR1 and APR2, although distant in sequence, are spatially adjacent in the ApoA-I crystal structure (Fig. 3c and 11a) and converge to form a steric zipper in the fibril core (Fig. 3b and 10b). In particular, the side chains from Val19 in APR1 and Thr56 in APR2 maintain proximity in both conformations (3.72 Å in the native, 4.31 Å in fibrils, Fig. S11), suggesting that this contact may nucleate the zipper interface. Additionally,

Thr54 may reorient inward, and together with Thr56, stabilize the APR1-APR2 steric zipper via coordination with two water molecules (Fig. S11b and S9h). The critical role of APR1 and APR2 in fibril formation of ApoA-I was first demonstrated *in vitro* by Mizuguchi et al., who showed that eliminating APR1 (residues 14–22) inhibited fibril formation of the G26R fragment spanning residues 1 to 83, whereas eliminating APR2 (residues 50–58) significantly reduced nucleation and fibril growth rates^47^.

Adding to this mechanism, our data reveal the presence of Pro66 in a *cis*-isomer conformation in both protofilaments of all determined structures (Fig. S9g). Proline isomerization, a unique type of post-translational conformational change, is known to play an important role in amyloid formation and proposed to act as a conformational switch that influences protein folding and refolding^48,49^. This conversion may occur either before (and potentially trigger) or after the disruption of ApoA-I’s α-helical secondary structure. In either case, the 180° flip of the Pro66 peptide bond would enable an additional hydrogen bond with Gln63 in the adjacent layer, further stabilizing the fibril core (Fig. S9g). These findings underscore the importance of APR exposure and proline isomerization of the native ApoA-I structure in AApoA-I amyloidosis.

## Conclusion

In summary, this study provides structural insights into *ex-vivo* AApoA-I fibrils obtained from multiple organs of patients carrying distinct mutations with varying clinical manifestations. The observed fibril heterogeneity, possibly driven by mutation location, may play a key factor in tissue tropism and disease outcome in hereditary AApoA-I amyloidosis. While only the L90P and R173P fibrils were structurally characterized, the tentative conclusions regarding G26R fibrils emphasize the need for further structural investigation. Our proteolysis data and APR analysis align with previous findings, reinforcing the central role of APRs in fibril formation. Furthermore, the discovery of Pro66 in a *cis*-isomer conformation in the fibril structure, but not in native ApoA-I, highlights the potential contribution of a conformational switch to amyloidogenesis. While our conclusions are based on a limited number of patient-derived samples, the trends observed support the need for broader investigation to fully elucidate the interplay between mutation, fibril structure, and clinical presentation.

## Methods

### Patients and tissue materials

We obtained human tissue samples from the laboratory of Dr. Merrill D. Benson at the Indiana University School of Medicine, with exemption from Internal Review Board oversight as they were anonymized. These specimens were postmortem frozen tissues from three patients diagnosed with AApoA-1 amyloidosis. Patient 1, a female carrying the G26R mutation, passed away at age 65 from renal failure; her kidney, spleen, and liver tissues were used in this study^50^. Patient 2, a female with the L90P mutation, died at age 60 from heart failure and also had cutaneous lesions. Heart tissue from this patient was used in this study^31^. Patient 3, a male with the R173P mutation, died at age 63 with cardiomyopathy, liver, and renal failure; his heart, kidney, spleen, and liver were used in this study^30^.

### Histology

Fresh-frozen human cardiac tissue was thawed overnight at 4°C and then fixed in 20x volume excess of 10% neutral buffered formalin for 48 hours at room temperature with agitation. After fixation, samples were transferred to 70% ethanol and paraffin processed according to established protocols^51^. Sections were generated from paraffin blocks for hematoxylin and eosin staining (H&E), Congo Red and Thioflavin-S staining. Regressive H&E was performed on a Sakura DRS-601 x,y,z robot utilizing Leica-Surgipath Selectech reagents according to Sheehan’s textbook methodology. Congo Red slides counterstained with hematoxylin were evaluated for pathologic amyloid aggregation under bright-field following their preparation. Color was imparted to amyloid by methods also described in Sheehan’s text utilizing 0.1% Congo Red in alcoholic-saline following sensitization of the slides with alkaline-alcoholic-saline (10% NaCl, 0.5% NaOH, and 80% EtOH). Thioflavin-S slides were evaluated for foci consistent with plaque amyloid and vascular amyloid, which fluoresce brilliantly when imaged with ultraviolet excitation (400-440nm) and long-pass emission (470 nm+). Thioflavin-S staining was accomplished utilizing methods described by Guntern, et.al.^52^. In brief, paraffin sections were brought to water and prepared for alcoholic Thioflavin-S impregnation by sequential oxidation with 0.25% w/v potassium permanganate, bleaching (1% potassium metabisulfite, 1% oxalic acid), peroxidation (1% hydrogen peroxide, 2% sodium hydroxide), and acidification (0.25% acetic acid), all with interceding water washes. Slides were then brought to 50% ethanol and incubated in alcoholic Thioflavin-S (0.004%). After seven minutes staining interval, slides were passed through alcoholic rinses to remove excess Thioflavin-S, dehydrated, cleared, and cover slips affixed with Cytoseal 60 permanent non-fluorescent mounting media (Epredia, Kalamazoo, MI).

### Extraction of AApoA-1 amyloid fibrils

AApoA-1 e*x-vivo* preparation of amyloid fibrils was obtained from the fresh-frozen human tissue as described earlier with slight modifications^32,53^. Briefly, ∼250 mg of frozen tissue was thawed at room temperature and minced into small pieces with a scalpel. The minced tissue was resuspended into 0.5 mL Tris-calcium buffer (20 mM Tris, 138 mM NaCl, 2 mM CaCl_2_, 0.1% NaN_3_, pH 8.0) and centrifuged for 5 min at 7000 × g and 4 °C. The pellet was washed in Tris-calcium buffer four additional times. After washing, the pellet was resuspended in 0.5 mL of 5 mg/mL collagenase solution (collagenase stock prepared in Tris-calcium buffer). Sample was spiked with complete protease inhibitor cocktail EDTA-free tablets (Roche) and incubated overnight at 37 °C with shaking at 700 rpm (Eppendorf ThermoMixer). The resuspension was centrifuged for 30 min at 7000 × g at 4 °C and the pellet was resuspended in 0.5 mL Tris–ethylenediaminetetraacetic acid (EDTA) buffer (20 mM Tris, 140 mM NaCl, 10 mM EDTA, 0.1% NaN_3_, pH 8.0). The suspension was centrifuged for 5 min at 7000 × g and 4 °C, and the washing step with Tris–EDTA was repeated nine additional times. After final washing, the pellet was resuspended in 150 μL ice-cold water supplemented with 5 mM EDTA. After a 15-minute incubation on ice, the sample was centrifuged for 5 min at 3100 × g and 4 °C. This extraction step was repeated three additional times and elutions were collected for further analysis.

### Negative-stained transmission electron microscopy

The extracted AApoA-1 amyloid fibril was confirmed by transmission electron microscopy. A 2.5 μL aliquot of the fibril sample was spotted onto a freshly glow-discharged carbon film 300 mesh copper grid (Electron Microscopy Sciences) and incubated for 1 min. Excess solution was gently blotted with filter paper. The grid was then stained with 2.5 µL of 2% uranyl acetate for 1 min, followed by gently blotting to remove excess stain. A second application of 2.5 μL uranyl acetate was immediately blotted, and the grid was air-dried vertically for at least 2 min. Specimens were imaged using an FEI Tecnai 12 electron microscope (Thermo Fisher Scientific) at an accelerating voltage of 120 kV.

### Mass Spectrometry (MS) sample preparation, data acquisition and analysis

For enzymatic MS analysis, 0.5 µg of extracted fibrils were dissolved in a tricine SDS sample buffer, boiled for 2 minutes at 85 °C, and run on a Novex™ 16% tris-tricine gel system using a Tricine SDS running buffer. Gel was stained with Coomassie dye, destained and the smear with AApoA-1 was cut from the gel. Sample was sent for MS analysis. Samples were digested overnight with trypsin (Pierce) following reduction and alkylation with DTT and iodoacetamide (Sigma–Aldrich). The samples then underwent solid-phase extraction cleanup with an Oasis HLB plate (Waters) and the resulting samples were injected into an Q Exactive HF mass spectrometer coupled to an Ultimate 3000 RSLC-Nano liquid chromatography system. Samples were dissolved in Buffer A (2% (v/v) ACN and 0.1% formic acid in water), injected onto a 75 µm i.d., 15-cm long EasySpray column (Thermo), and eluted with a gradient from 0-28% buffer B over 90 min. Buffer B contained 80% (v/v) ACN, 10% (v/v) trifluoroethanol, and 0.1% formic acid in water. The mass spectrometer operated in positive ion mode with a source voltage of 2.5 kV and an ion transfer tube temperature of 300 °C. MS scans were acquired at 120,000 resolution in the Orbitrap and up to 20 MS/MS spectra were obtained in the ion trap for each full spectrum acquired using higher-energy collisional dissociation (HCD) for ions with charges 2-8. Dynamic exclusion was set for 20 s after an ion was selected for fragmentation. For identification of AApoA-1 R173P mutation, Glu-C enzyme (Promega) was used instead of trypsin, with no other steps in the protocol altered.

Raw MS data files were analyzed using Proteome Discoverer v3.0 SP1 (Thermo), with peptide identification performed using a semitryptic search with Sequest HT against the human reviewed protein database from UniProt. Fragment and precursor tolerances of 10 ppm and 0.02 Da were specified, and three missed cleavages were allowed. Carbamidomethylation of Cys was set as a fixed modification, with oxidation of Met set as a variable modification. The false-discovery rate (FDR) cutoff was 1% for all peptides.

For Intact protein mass analysis, 20 µL (∼5 µg) of extracted AApoA-1 fibrils were spin at 21000 x g for 1 h at 4 °C. The 15 µL of supernatant was removed and 5 µL of 8M GuHCl was added to dissociate the fibrils. Mixture was incubated at 37 °C for 1 h with continuous shaking at 1000 rpm (Eppendorf ThermoMixer). Then, an additional 5 µL of 8M GuHCl was added and incubated at 37 °C for 1 at with continuous shaking at 1000 rpm. Finally, 5 µL of MilliQ water containing reducing agent (DTT) was added. Samples were desalted and analyzed by LC/MS, using a Sciex X500B QTOF mass spectrometer coupled to an Agilent 1290 Infinity II HPLC. Samples were injected onto a POROS R1 reverse-phase column (2.1 × 30 mm, 20 µm particle size, 4000 Å pore size) and desalted. The mobile phase flow rate was 300 uL/min and the gradient was as follows: 0-3 min: 0% B, 3-4 min: 0-15% B, 4-16 min: 15-55% B, 16-16.1 min: 55-80% B, 16.1-18 min: 80% B. The column was then re-equilibrated at initial conditions prior to the subsequent injection. Buffer A contained 0.1% formic acid in water and buffer B contained 0.1% formic acid in acetonitrile.

The mass spectrometer was controlled by Sciex OS v.3.3.1.43 using the following settings: Ion source gas 1 30 psi, ion source gas 2 30 psi, curtain gas 35, CAD gas 7, temperature 300 °C, spray voltage 5500 V, declustering potential 135 V, collision energy 10 V. Data was acquired from 400-2000 Da with a 0.5 s accumulation time and 4 time bins summed. The acquired mass spectra for the proteins of interest were deconvoluted using Bio Tool Kit within SciexOS in order to obtain the molecular weights. Peaks were deconvoluted over the entire mass range of the mass spectra, with an output mass range of 7000-9000 Da, using low input spectrum isotope resolution.

The mass spectrometry data have been deposited to MassIVE data repository (a member of ProteomeXchange), with accession number MSV000098187.

### Cryo-EM sample preparation and data collection

Freshly extracted fibril samples were applied to glow-discharged R1.2/1.3, 300-mesh copper grids (Quantifoil), blotted with filter paper to remove excess sample, and plunge-frozen in liquid ethane using a Vitrobot Mark IV (FEI/Thermo Fisher Scientific). Cryo-EM samples were screened on Talos Arctica at the Cryo-Electron Microscopy Facility (CEMF) at The University of Texas Southwestern Medical Center (UTSW). Images were collected using a 300 keV Titan Krios microscope (FEI/Thermo Fisher Scientific) with Falcon4i detector operated at slit width of 10 eV at the Stanford-SLAC Cryo-EM Center (S^2^C^2^). Further details are provided in Supplementary Table 2.

### Helical reconstruction

The raw movie frames were gain-corrected, aligned, motion-corrected and dose-weighted using RELION’s own implemented motion correction program^54,55^. Contrast transfer function (CTF) estimation was performed using CTFFIND 4.1^56^. All steps of helical reconstruction, three-dimensional (3D) refinement, and post-process were carried out using RELION 4.0 and RELION 5.0 beta^57,58^. The filaments were picked automatically using Topaz in RELION 4.0^59,60^. Particles were extracted using box size of 1024 and 300 or 256 pixels with an inter-box distance of 3 asymmetrical units at helical rise of 4.75 Å. 2D classification of 1024-pixel particles was only used to estimate the fibril crossover distance. 2D classifications of 300-pixel particles were used to select suitable particles for further processing. An initial 3D reference model were generated from a subset of 2D class averages using relion_helix_inimodel2d program, as previously described^61^. Fibril helix was assumed left-handed for 3D reconstruction. 3D auto-refinement was followed by 3D classification without image alignment to filter out suboptimal classes, after which selected particles underwent further 3D auto-refinement. Subsequent 3D auto refinements with optimization of helical twist and rise were carried out once the estimated resolution of the map reached beyond 4.75 Å. Multiple rounds of CTF refinement, 3D classification (without image alignment), auto-refinement, and post-processing were performed sequentially to reach the highest possible resolution. As the optimized rise of all three structures were approximately 4.91 Å, we performed the final post-processing job in which we calibrated the pixel size to 0.923 Å/px (from 0.954 Å/px), corresponding to the rise of 4.75 Å. Overall resolutions were calculated from Fourier shell correlations at 0.143 between two independently refined half-maps using a soft-edged solvent mask^62^. Local resolutions were estimated using RELION’s Local Resolution tool. Additional processing details are available in Supplementary Table 2.

### Atomic model building and refinement

We used the automated machine-learning ModelAngelo approach with minor modifications to obtain an initial atomic model^63^. First, COOT v0.9.8.1 was used to build the peptide backbone of a single fibril layer featuring all alanine residues^64^. We then created a new density map of one fibril layer in ChimeraX v1.8 using the command ‘*vol zone #1 near #2 range 3 new true*’, where #1 refers to the post-processed fibril density map and #2 is the peptide backbone model^65^. This new map was then entered ModelAngelo, both with and without the primary ApoA-1 sequence, to obtain the initial atomic models of AApoA-1 fibrils. Using COOT, we made residue modifications and real-space refinements to finalize the model. Further refinement was carried out using ‘*phenix*.*real_space_refine*’ from PHENIX 1.20^66^. ChimeraX v1.8 was used for molecular graphics and structural analysis^67^. All refinement statistics are summarized in Supplementary Table 2.

### Stabilization energy calculation

Solvation energy per residue was determined by summing the products of each atom’s buried surface area and its atomic solvation energy^68^. The total chain energy was calculated by summing the residue energies, and each residue was assigned a distinct color in the solvation energy map to visualize variations in energy distribution.

Link to download the program: https://doi.org/10.5281/zenodo.6321286.

### Aggregation-prone regions analysis

Aggregation-prone regions (APRs) were predicted using four common prediction models: TANGO^69-71^, WALTZ^72^, Aggrescan^73^, and AmylPred^74^. For each ApoA1 sequence variant (WT, G26R, L90P, and R173P), residues predicted as APRs by a model (“hit”) were given a score of 1, while residues not part of an APR were given a score of 0. This scoring process was repeated across all four models, yielding a per-residue score based on the sum of these hits. For each model, APR criteria were as follows: in TANGO, any 5-residue or longer with a score ≥ 5.0; in WALTZ, residues with a score > 0; in Aggrescan, 5-residue stretches with a score ≥ 0.02; and in AmylPred, a score of 3 or more. Total scores were recorded in a PDB file as B-factors for visualizing the heatmap model, generated in ChimeraX v1.8^67^.

## Data and Materials Availability

Mass spectrometry data have been deposited to MassIVE database (a member of ProteomeXchange) under accession code MSV000098187. Cryo-EM maps and their models are deposition codes are to be updated.

## Figure Panels

Figure panels were created with ChimeraX v1.8 and Adobe Illustrator.

## Acknowledgments

Special thanks to the patients and families who generously donated tissues and the University of Indiana as the source of material.

We thank all members of the Structural Biology Laboratory (SBL), the UTSW Cryo-Electron Microscopy Facility and the UTSW Electron Microscopy Core Facility for instrumentation and technical support. SBL and CEMF at UTSW are partially supported by grant RP220582 from the Cancer Prevention & Research Institute of Texas (CPRIT). We thank the UTSW Proteomics Core for assistance with proteomics analysis and Histo Pathology Core for assistance with histological analysis.

We special thank the national cryo-EM facilities Stanford-SLAC (project CA172) for instrumentation, technical support, and data collection. Cryo-EM Data collection of all samples from this work was performed at the Stanford-SLAC Cryo-EM Center (S2C2), which is supported by the National Institute of General Medical Sciences (1R24GM154186). The content is solely the responsibility of the authors and does not necessarily represent the official views of the National Institutes of Health. The authors would also like to thank the following S2C2 personnel for their invaluable support and assistance: Lisa B. Dunn, Ian M. Fries, Dr. Alexandre Cassago and Dr. Patrick Mitchell.

This research was supported in part by the computational resources provided by the BioHPC supercomputing facility located in the Lyda Hill Department of Bioinformatics, UT Southwestern Medical Center, TX. URL: https://portal.biohpc.swmed.edu.

Molecular graphics and analyses performed with UCSF ChimeraX, developed by the Resource for Biocomputing, Visualization, and Informatics at the University of California, San Francisco, with support from National Institutes of Health R01-GM129325 and the Office of Cyber Infrastructure and Computational Biology, National Institute of Allergy and Infectious Diseases.

We used ChatGPT (OpenAI, 2025) to help edit and improve the clarity of the manuscript. The authors reviewed and verified all AI-assisted content.

## Funding

American Heart Association (Career Development Award 847236).

National Institutes of Health, National Heart, Lung, and Blood Institute (New Innovator Award DP2-HL163810).

Welch Foundation (Research Award I-2121-20220331).

UTSW Endowment (Distinguished Researcher Award from President’s Research Council and start-up funds).

## Author contributions

Conceptualization: L.S., V.S., B.A.N.

Methodology: L.S., V.S., B.A.N.

Investigation: V.S., B.A.N., P.B., Y.A., P.S., M.P., M.C.F.R., L.V., A.L., B.E., C.L., B.K.B, L.S.

Visualization: B.A.N., L.S.

Funding acquisition: L.S.

Project administration L.S., V.S., B.A.N

Supervision: B.A.N, L.S.

Writing – original draft: B.A.N., M.C.F.R., V.S.

Writing – review & editing: M.C.F.R., V.S., B.A.N., L.S.

## Competing interests

L.S. reports research funding from NHLBI, Welch Foundation, UTSW, and AstraZeneca. L.S. also reports advisory board, speaker, and consulting fees from Alexion, Pfizer, Attralus, Intellia, and AmyGo.

B.A.N. also reports advisory board, speaker, and consulting fees from Amygo.

