## Supplementary information for "Structural variability of apolipoprotein A-I amyloid fibrils across organs, mutations, and clinical presentations, revealed by cryo-EM"

**Affiliations:**
[1]Center for Alzheimer's and Neurodegenerative Diseases, Department of Biophysics, Peter O'Donnell Jr Brain Institute, University of Texas Southwestern Medical Center (UTSW), Dallas, TX, USA.
[2]SciKonnect and BioPatriKa, India.
[3]Department of Biochemistry, University of Texas Southwestern Medical Center, Dallas, TX, USA
[4]Department of Pathology, University of Texas Southwestern Medical Center (UTSW), Dallas, TX, USA.
[5]Department of Ophthalmology, University of Texas Southwestern Medical Center (UTSW), Dallas, TX, USA.
[6]Department of Internal Medicine, University of Texas Southwestern Medical Center (UTSW), Dallas, TX, USA.
[7]Department of Pathology and Laboratory Medicine, Indiana University School of Medicine, Indianapolis, IN, USA.

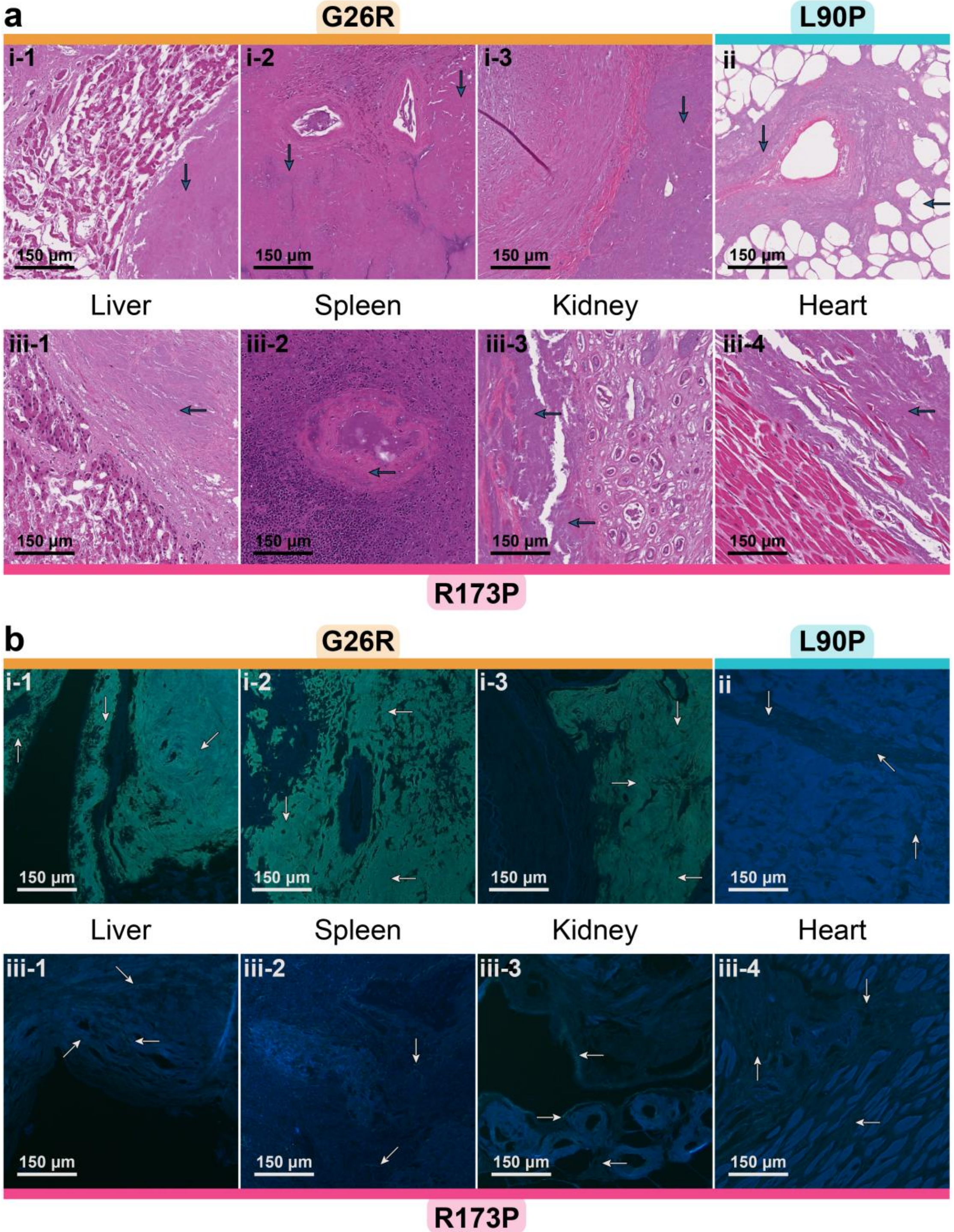


**Supplementary Figure 1. Histological analysis of AApoA-I amyloid deposits** of various tissue samples from three patients: G26R, L90P and R173P using (**a**) hematoxylin and eosin, and (**b**) Thioflavin S stains. Arrows indicate amyloid deposits. Cross sections show amyloid in the liver, spleen and kidney of G26R (**i1-3**), the heart of L90P (**ii**) and the liver, spleen, kidney and heart of R173P (**iii1-4**). Scale bar: 150 µm.

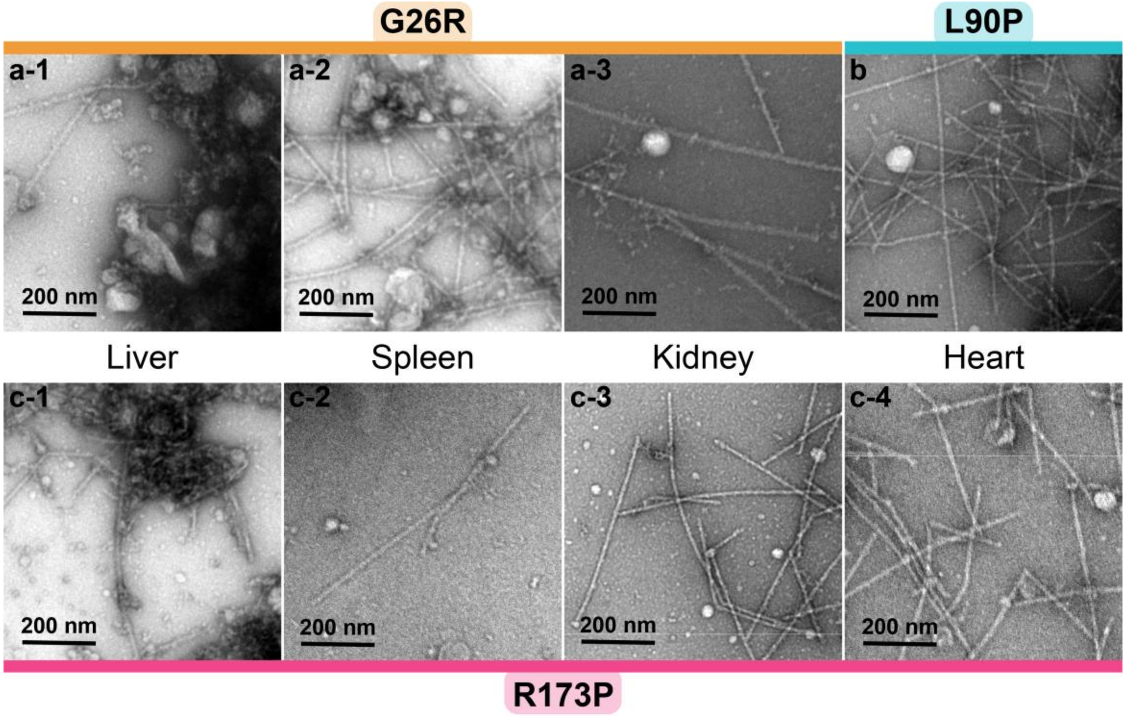


**Supplementary Figure 2. Negatively stained transmission electron microscopy (TEM) images of AApoA-I fibrils** extracted from various tissue samples in three patients: G26R, L90P and R173P. TEM images of fibrils extracted from the liver, spleen and kidney of G26R (**a1-3**); fibrils from the heart of L90P (**b**); and the liver, spleen, kidney and heart of R173P (**c1-4**). Scale bar: 200 μm.

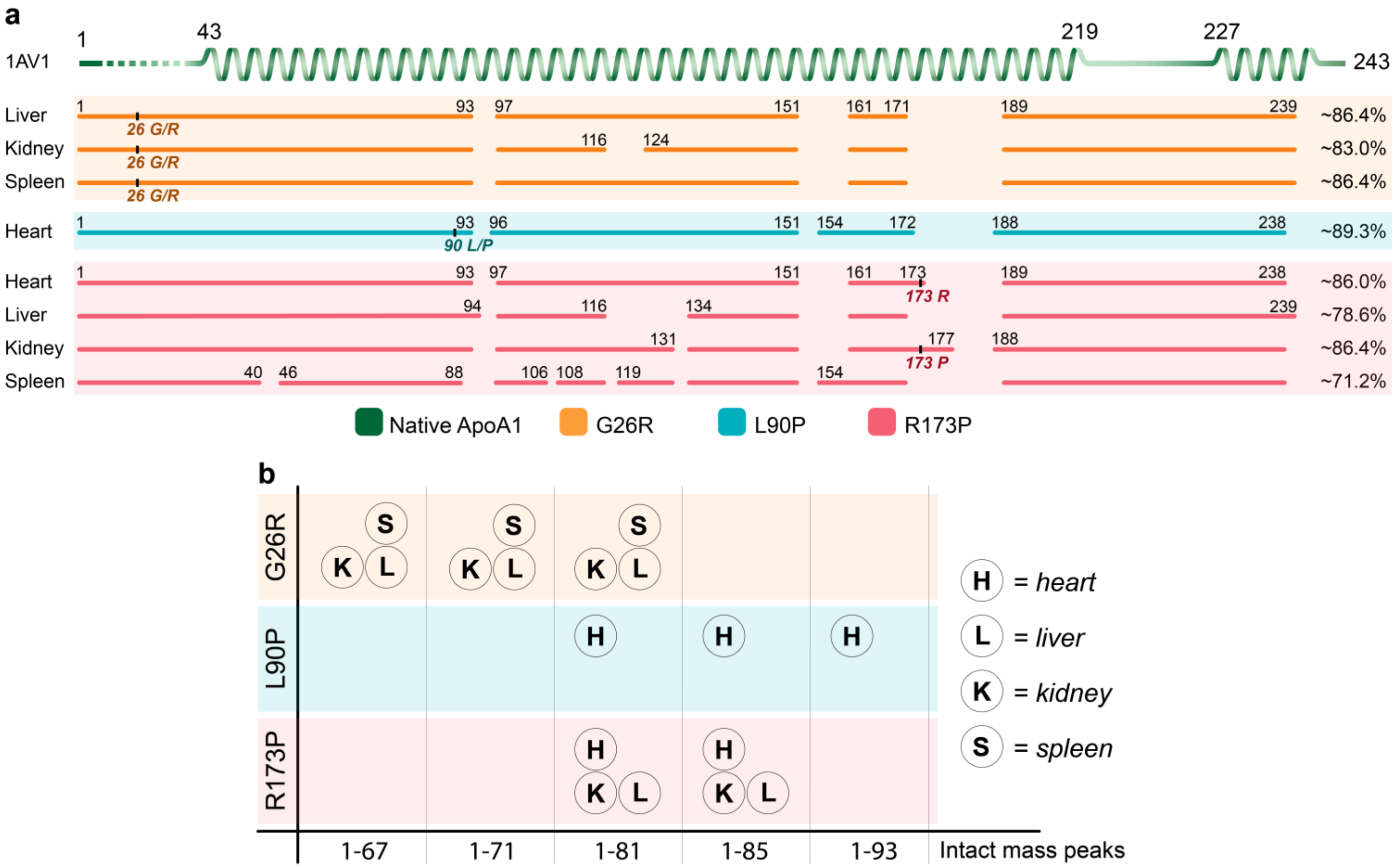


**Supplementary Figure 3. Mass spectrometry analysis of fibrils extracted from three AApoA-I patients: G26R, L90P and R173P. (a)** Tryptic digest analysis of all fibril samples and GluC analysis for only R173P sample, with the native sequence shown in green (α-helix structure indicated, pdb: 1AV1). Fibril fragments are color-coded by patients: G26R (orange), L90P (blue), and R173P (pink). Detected residues are shown as continuous lines with the corresponding residue positions indicated above each sequence. Variant-specific residues (wild-type and mutant) are noted at their respective positions. Total sequence coverage for each sample is provided at the end. (**b**) Intact mass analysis identifying peaks corresponding to AApoA-I fragments, with fibril samples color-coded as in (a). H, L, K, and S represent fibrils from the heart, liver, kidney, and spleen, respectively.

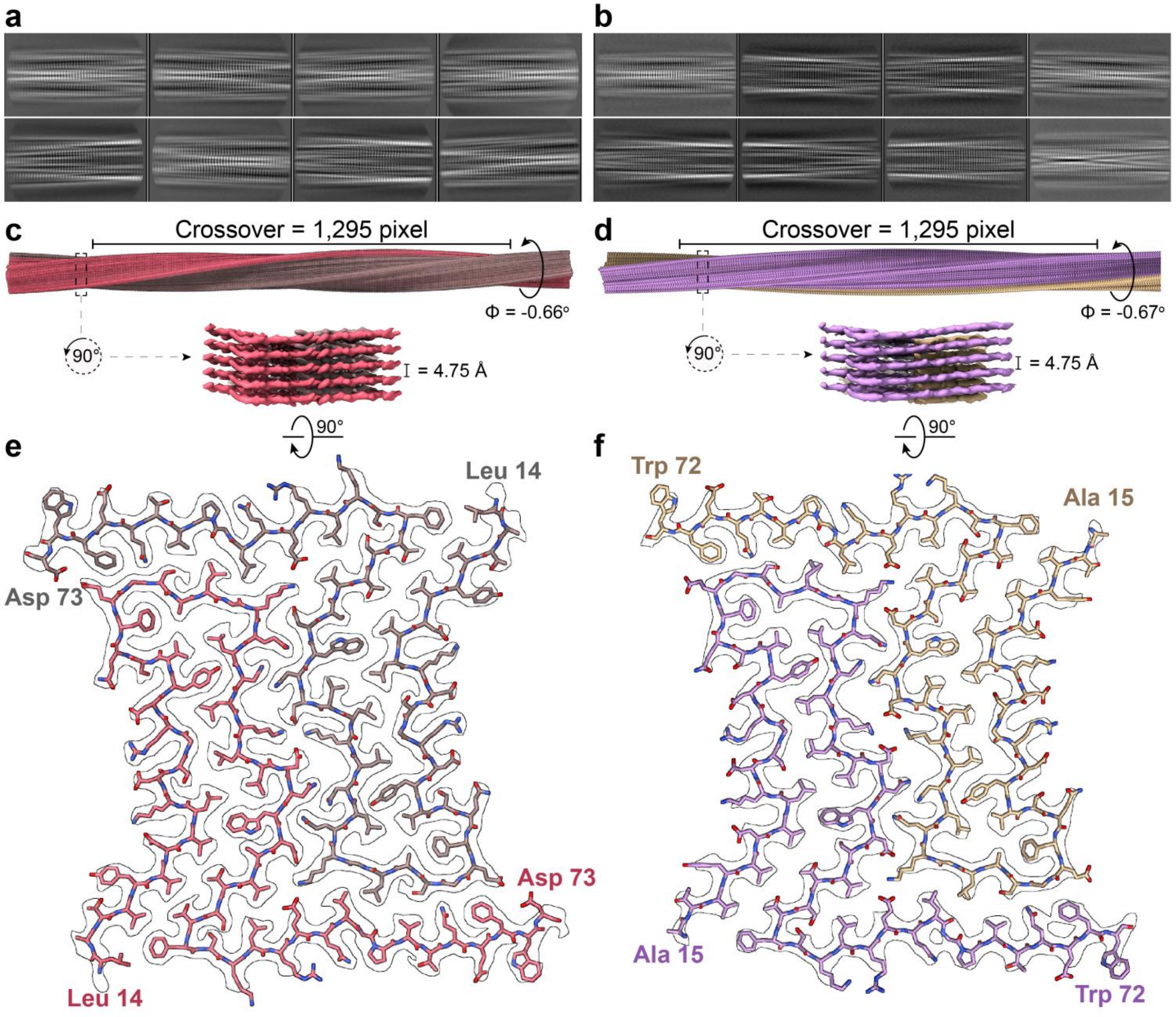


**Supplementary Figure 4. Helical reconstructions of AApoA-I fibrils from R173P samples.** (a) and (b) 2D class averages of R173P fibrils from the heart and kidney, respectively. (c) and (d) The reconstructed R173P fibrils from the heart and kidney, respectively, containing 360 layers and the magnified view of a subset of 5 fibril layers. (e) and (f) Top view of a single fibril layer from R173P fibrils (heart and kidney, respectively) with a model consisting of two protofilaments spanning residues Leu 14 or Ala 15 to Asp 73 or Trp 72.

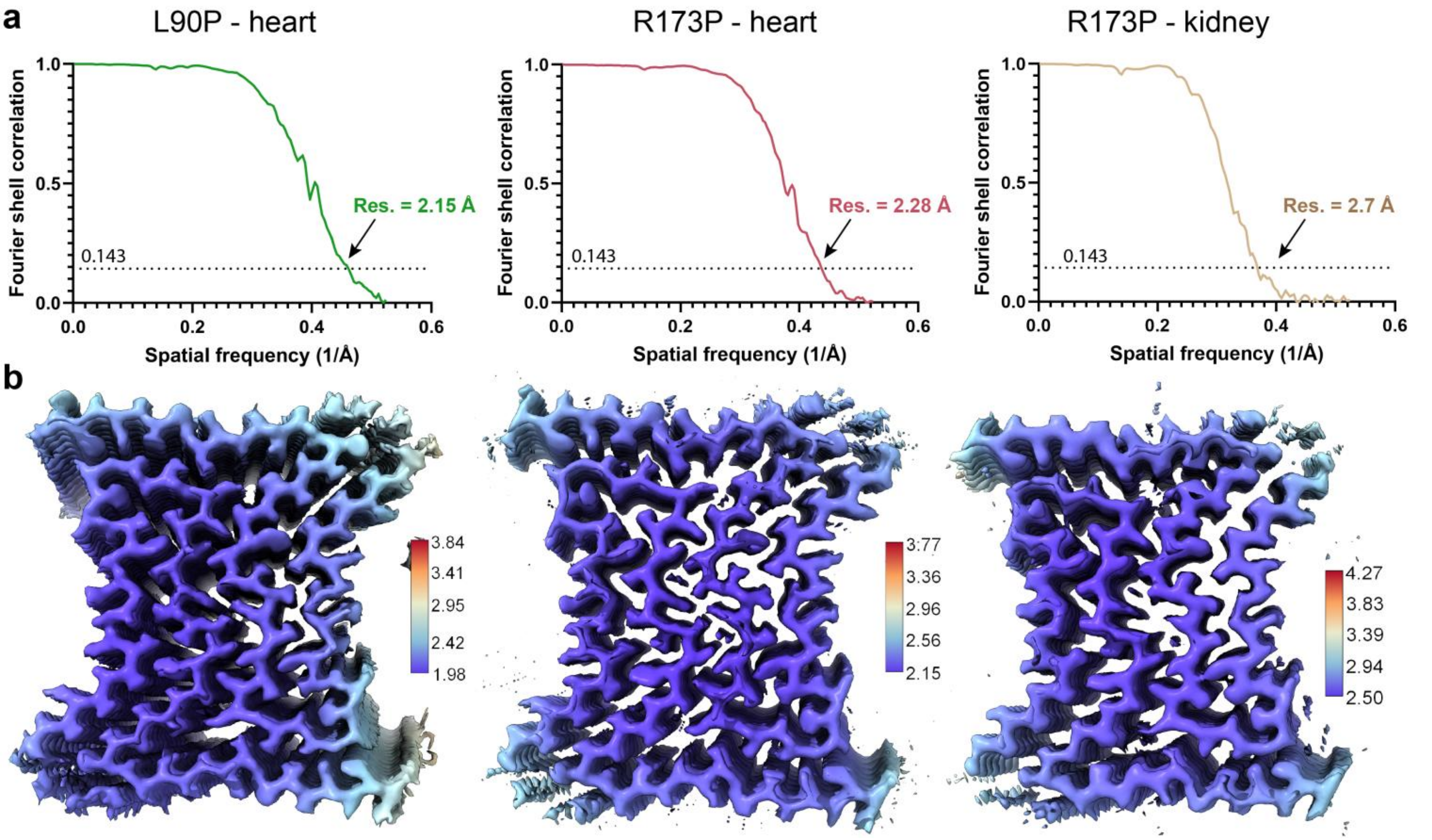


**Supplementary Figure 5. FSC curves and local resolution maps of cryo-EM structures from L90P heart, R173P heart and kidney**. **(a-c)** Evaluation of the resolution of cryo-EM maps by Fourier shell correlation (FSC) curves of two independently refined half-maps from these samples. **(d-f)** Local resolution estimation for 3D reconstructions of AApoA-I fibrils. Resolution scale in Angstroms.

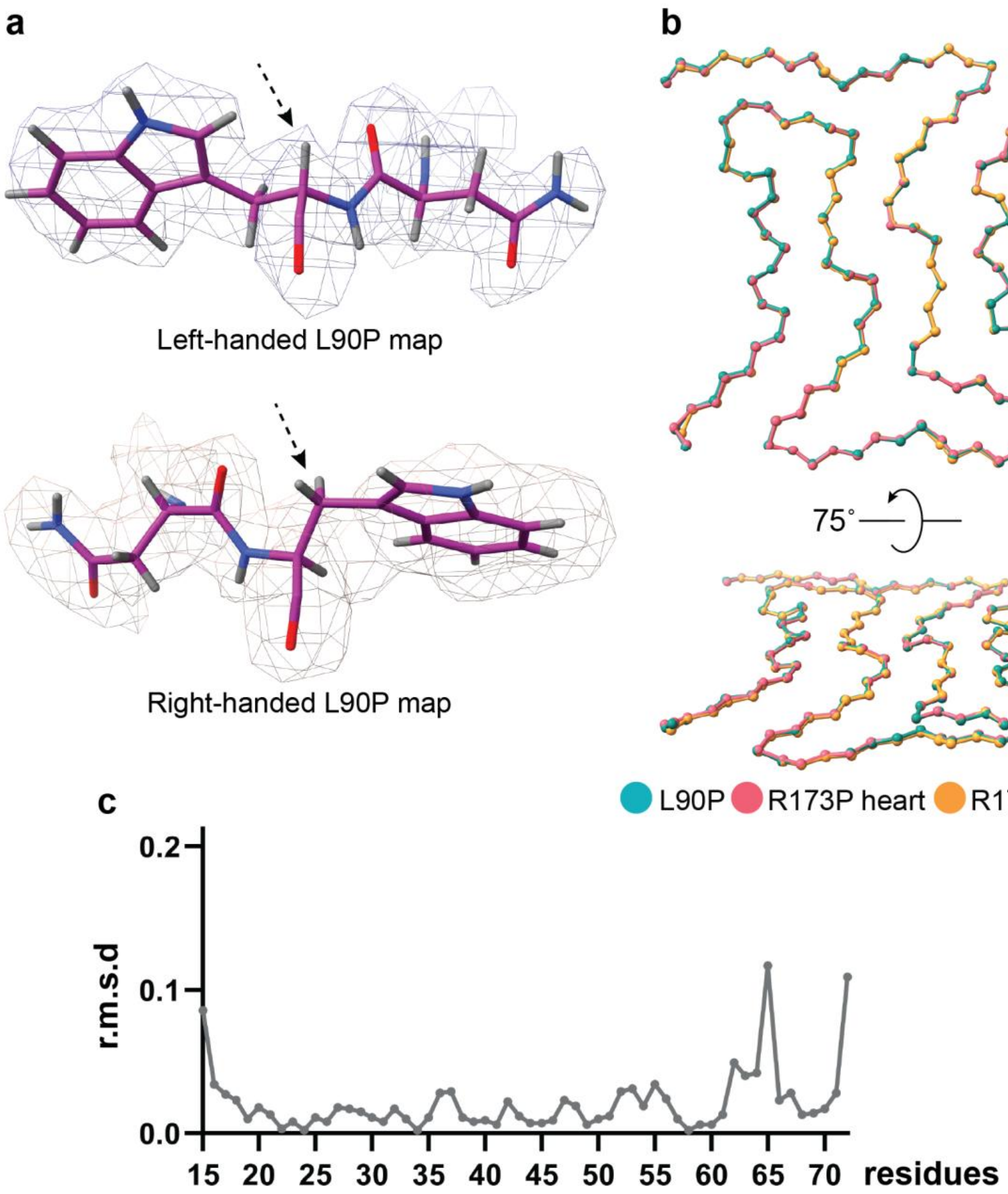


**Supplementary Figure 6. Handedness of AApoA-I fibril maps and their structural alignment. (a)** representative L-amino acids demonstrate a better fit to the left-handed L90P map compared to the right-handed map. **(b)** Top view and tilted view of structural alignment of the Cα backbones from L90P fibrils (teal) and R173P fibrils from heart (light red) and kidney (yellow) samples. **(c)** Comparison of Cα displacement among these three structures, estimated using GESAMT, shows that the majority of displacements are less than 0.1 Å compared to the consensus sequence of the three fibril structures, revealing highly structural similarity.

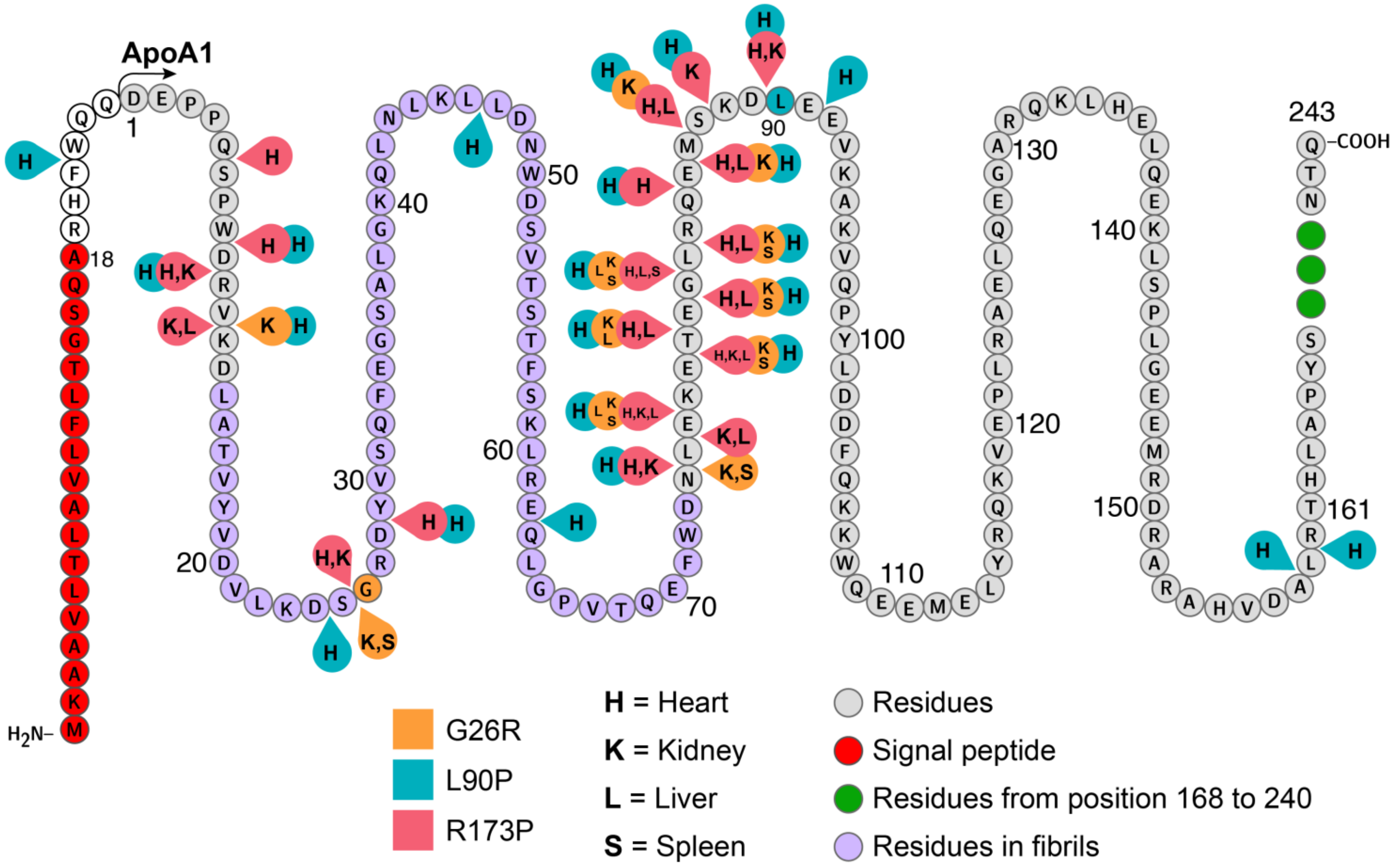


**Supplementary Figure 7. Combined proteolytic sites from all three variants analyzed by mass-spectrometry**. The signal peptide is highlighted in red. Regions corresponding to G26R, L90P and R173P variants are shown in light orange, blue, and salmon pink, respectively. The fibril core is depicted in purple, and residues from positions 168 to 240 are shown in green. Proteolytic sites identified in the heart, kidney, liver and spleen are denoted by H, K, L, and S, respectively.

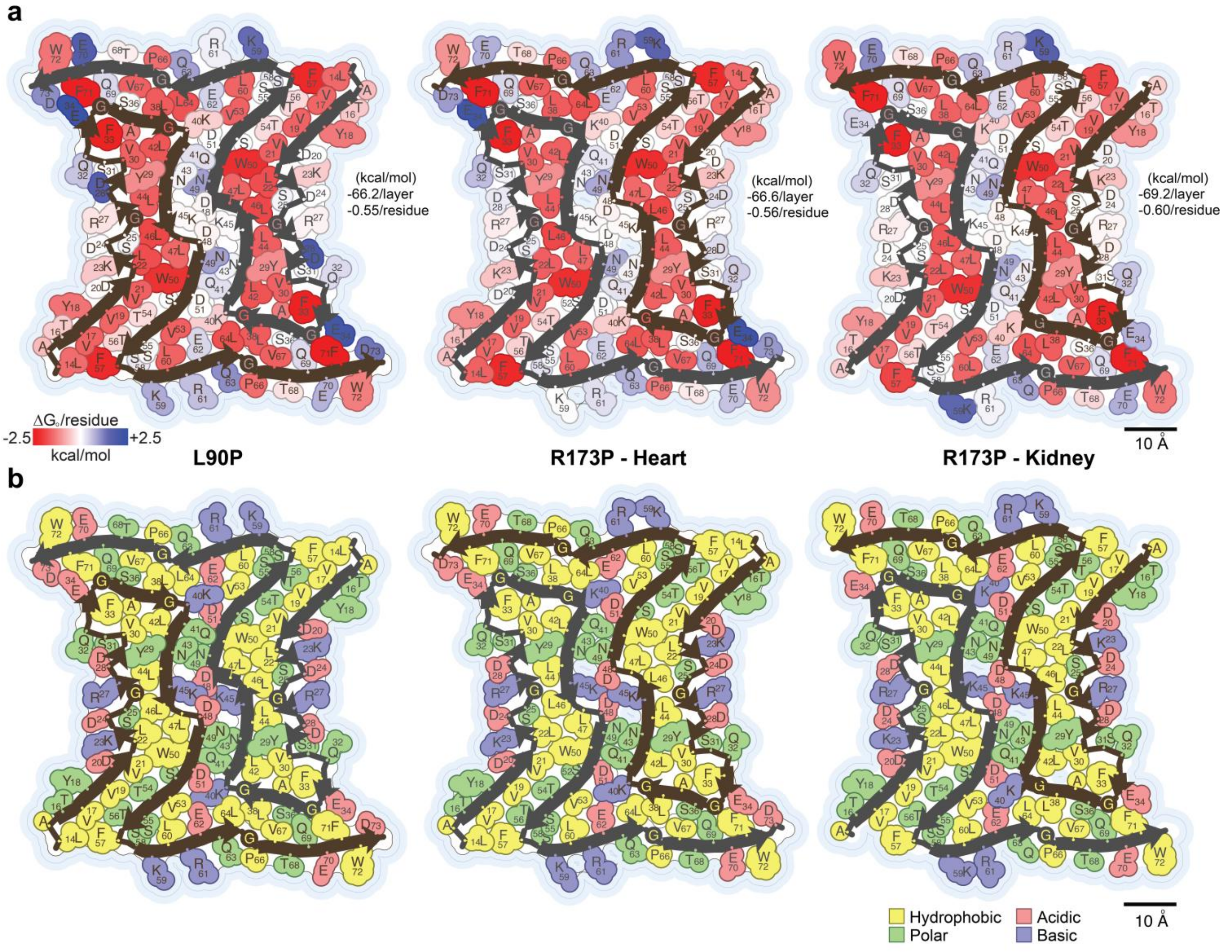


**Supplementary Figure 8**. **Solvation energy and fibril composition profile of L90P and R173P amyloidosis**. **(a)** Residue solvation energies profiles for fibril structures from the L90P heart, R173P heart, and R173P kidney, with stabilization energies ranging from favorable (red, -2.5 kcal/mol) to unfavorable (blue, 2.5 kcal/mol). Scale: 10 Å. **(b)** Residue composition profiles of fibril structures from the same samples, with residue color-coded as hydrophobic (yellow), polar (green), acidic (red), and basic (blue). Scale, 10 Å.

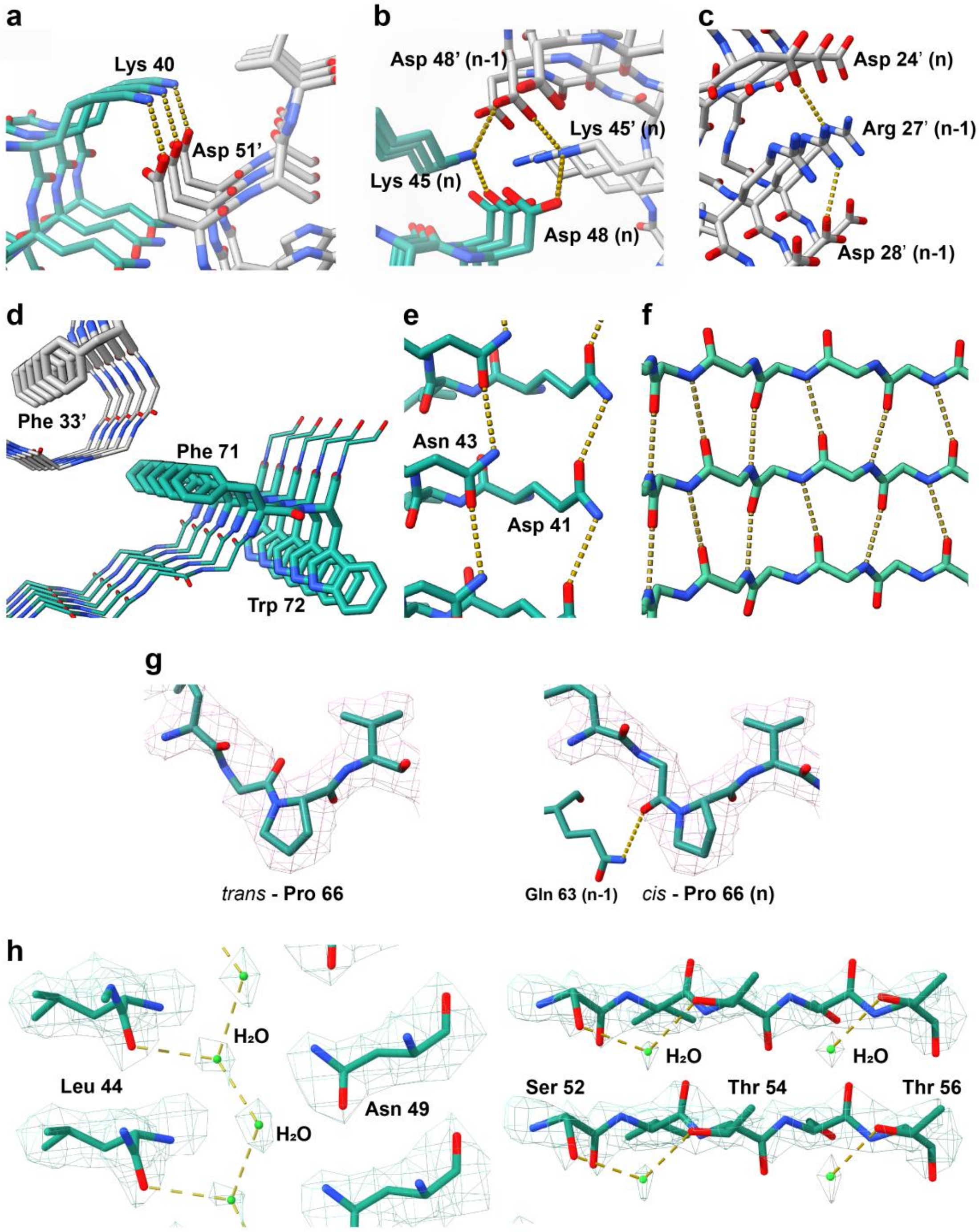


**Supplementary Figure 9. Molecular interactions stabilize AApoA-I fibril structure, the cis-isomer Pro 66 and densities of likely water molecules**. (**a**) Salt bridges (approximately 2.7 Å) formed between Lys 40 on layer n and Asp 51' on layer n-1 of the adjacent protofilament. (**b**) Salt bridges (approximately 2.5 Å) formed between Lys 45 on layer n and Asp 48' and Asp 48 on layer n-1. (**c**) Salt bridges formed within the same layer (n-1) between Arg 27' and Asp 28, and between Arg 27' of layer n-1 and Asp 24' of layer n. (**d**) π-π interactions from stacking side chains of aromatic residues. (**e, f**) H-bonds formed by layer-stacking aspartate and glutamate side chains, and peptide backbone, respectively. (**g**) Density map and the model of trans-Pro 66 (on the left) and cis-Pro 66 (on the right) with the additional hydrogen bond with Gln 63 (n-1). (**h**) Inter-layer water molecules forming hydrogen bonds with each other and with Leu 44 along the fibril axis (on the left) and intra-layer water molecules forming hydrogen bonds with Ser 52 and Thr 54 (on the right).

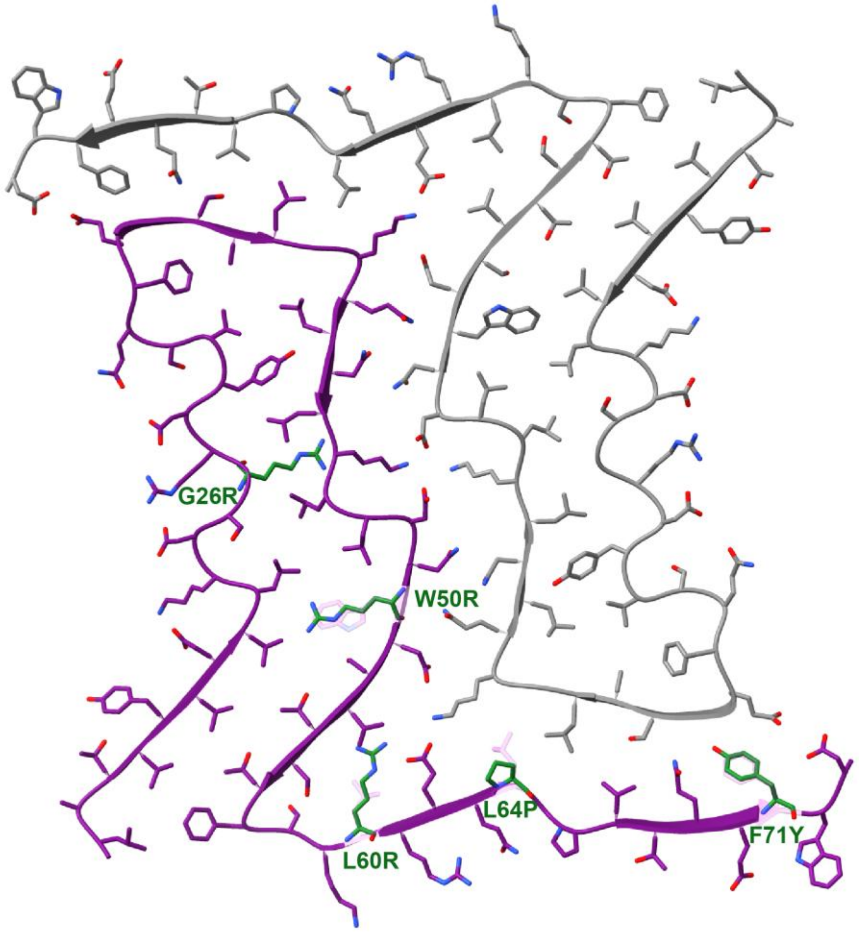


**Supplement Figure 10. Schematic representation illustrates how the introduction of N-terminal mutations into the fibrils core of L90P and R173P samples may influence fibril structures**. These mutations could potentially alter the structural integrity and stability of the fibrils due to changes of side chain properties such as bulkiness, charge, or rigidity. Mutated residues are colored green and native residues are dimmed in the background.

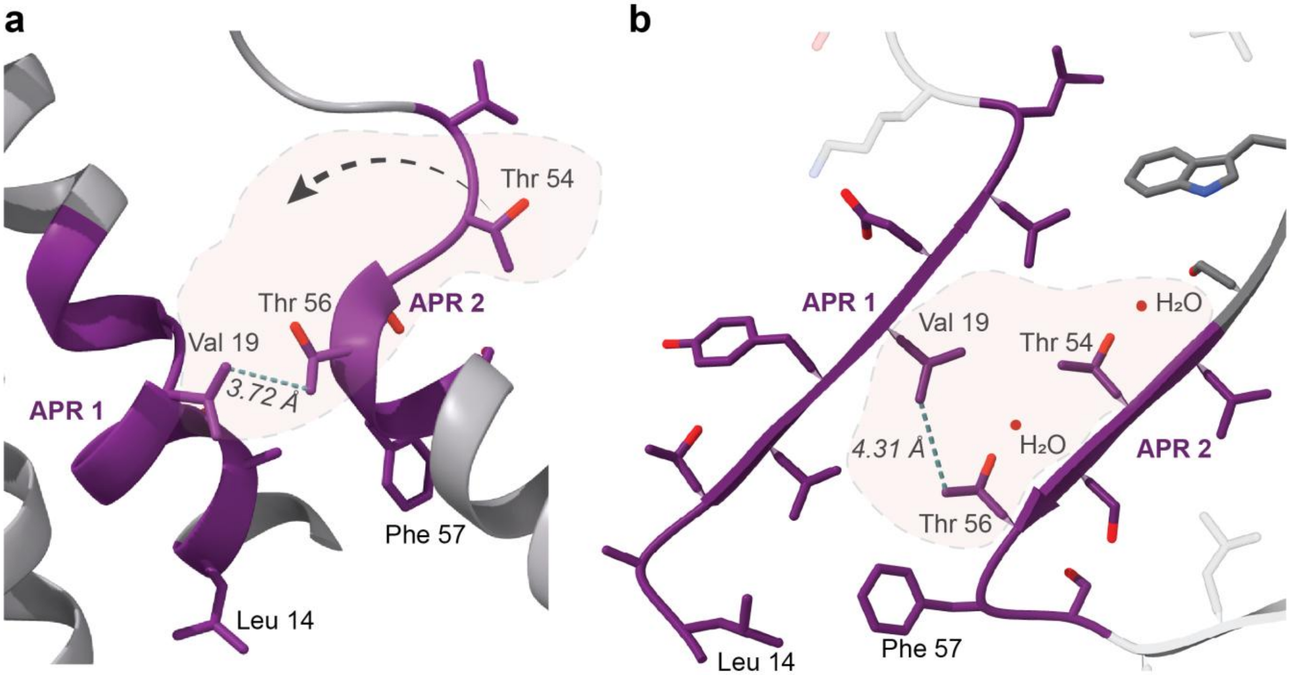


**Supplementary Figure 11: The two APR (1 and 2, in purple) of ApoA-I are spatially adjacent in both the native and fibril structure of the L90P, R173P samples**. **(a)** In the native crystal structure (PDB: 3R2P), APR1 and APR2 are positioned near each other, with Val 19 on APR1 located 3.72 Å from Thr 56 on APR2. **(b)** In the fibril structures, APR1 and APR2 remain close, with Val 19 on APR1 being 4.31 Å from Thr 56 on APR2.

**Supplement Table 1: ApoA-I variants and the corresponding clinical manifestations [1-4].**

| Variants | Amyloid in kidney biopsy | Major organ involvement |
|---|---|---|
| D20Y | No | Esophageal |
| G26R | Yes | Renal failure |
| E34K | Yes | Renal failure |
| W50R | Yes | Renal failure |
| L60R | Yes | Renal failure |
| ∆60-71/60V61T | Yes | Hepatic failure |
| L64P | Yes | Renal failure |
| ∆70-72 | Yes | Renal failure |
| F71Y | n/r | Hepatic involvement |
| N74fs | n/r | Renal involvement |
| L75P | Yes | Renal failure |
| L90P | n/r | Cardiac failure |
| ∆K107 | n/r | Cardiomyopathy |
| A154fs | Yes | n/r |
| H155Mfs | Yes | Renal failure |
| L170P | n/r | Laryngeal |
| Q172P | n/r | Cardiac failure |
| R173P | n/r | Cardiomyopathy |
| L174S | n/r | Cardiac failure |
| A175P | n/r | Laryngeal |
| L178(H/R) | n/r | Cardiomyopathy |

n/r = not report. Purple represents the fibril core. Light blue represents the C-terminal outside of fibril core.

### Supplementary Data Table 2: Cryo-EM data collection, refinement and validation statistics

| | AApoA-I-G26R kidney | AApoA-I-G26R spleen | AApoA-I-L90P heart (EMDB-xxxx) (PDB xxxx) | AApoA-I-R173P heart (EMDB-xxxx) (PDB xxxx) | AApoA-I-R173P kidney (EMDB-xxxx) (PDB xxxx) |
|---|---|---|---|---|---|
| **Data collection and processing** | | | | | |
| Microscope | Titan Krios G3i | Titan Krios G3i | Titan Krios G3i | Titan Krios G3i | Titan Krios G3i |
| Detector | Falcon 4i | Falcon 4i | Falcon 4i | Falcon 4i | Falcon 4i |
| Magnification | 130,000 | 130,000 | 130,000 | 130,000 | 130,000 |
| Voltage (kV) | 300 | 300 | 300 | 300 | 300 |
| Exposure time (s) | 4.05 | 4.20 | 4.16 | 4.19 | 3.54 |
| Electron exposure (e-/Å$^2$) | 40 | 40 | 40 | 40 | 40 |
| Defocus range (μm) | -0.8 to -2.6 | -0.8 to -2.6 | -0.8 to -2.6 | -0.8 to -2.6 | -0.8 to -2.6 |
| Pixel size (Å/px)) | 0.954 | 0.954 | 0.954 | 0.954 | 0.954 |
| **Reconstruction** | | | | | |
| Micrographs | 16,361 | 13,044 | 12,645 | 16,496 | 5,298 |
| Box size (pixel) | 1,024 | 1,024 | 256 | 300 | 300 |
| Initial segments (no.) | 1,334,546 | 632,144 | 5,158,231 | 6,860,279 | 967,103 |
| Final segments (no.) | n/a | n/a | 118,342 | 165,734 | 133,438 |
| Symmetry imposed | n/a | n/a | C1 | C1 | C1 |
| Helical rise (Å) | n/a | n/a | 4.75 | 4.75 | 4.75 |
| Helical twist (°) | n/a | n/a | -0.68 | -0.66 | -0.67 |
| Map resolution (Å) (FSC = 0.143) | n/a | n/a | 2.15 | 2.29 | 2.73 |
| Map sharpening *B* factor (Å$^2$) | n/a | n/a | -66.47 | -77.32 | -96.14 |
| **Model** | | | | | |
| Initial model used (PDB code) | n/a | n/a | *De novo* | *De novo* | *De novo* |
| Model resolution (Å) (FSC = 0.143) | n/a | n/a | 2.15 | 2.29 | 2.72 |
| **Model composition** | | | | | |
| Chains | n/a | n/a | 10 | 10 | 10 |
| Non-hydrogen atoms | n/a | n/a | 4810 | 4,810 | 4,650 |
| Protein residues | n/a | n/a | 600 | 600 | 580 |
| **R.M.S. deviations** | | | | | |
| Bond lengths (Å) | n/a | n/a | 0.004 | 0.003 | 0.004 |
| Bond angles (°) | n/a | n/a | 0.728 | 0.672 | 0.683 |
| **Validation** | | | | | |
| MolProbity score | n/a | n/a | 2.01 | 1.85 | 1.81 |
| Clashscore | n/a | n/a | 11.28 | 7.38 | 11.55 |
| Poor rotamers (%) | n/a | n/a | 0 | 0 | 0 |
| **Ramachandran plot** | | | | | |
| Favored (%) | n/a | n/a | 93.10 | 93.10 | 96.43 |
| Allowed (%) | n/a | n/a | 6.90 | 6.90 | 3.57 |
| Disallowed (%) | n/a | n/a | 0 | 0 | 0 |
